# Identification of common key genes and pathways between Covid-19 and lung cancer by using protein-protein interaction network analysis

**DOI:** 10.1101/2021.02.16.431364

**Authors:** Kang Soon Nan, Kalimuthu Karuppanan, Suresh Kumar

## Abstract

COVID-19 is indeed an infection that is caused by a recently found coronavirus group, a type of virus proven to cause human respiratory diseases. The high mortality rate was observed in patients who had pre-existing health conditions like cancer. However, the molecular mechanism of SARS-CoV-2 infection in lung cancer patients was not discovered yet at the pathway level. This study was about determining the common key genes of COVID-19 and lung cancer through network analysis. The hub genes associated with COVID-19 and lung cancer were identified through Protein-Protein interaction analysis. The hub genes are ALB, CXCL8, FGF2, IL6, INS, MMP2, MMP9, PTGS2, STAT3 and VEGFA. Through gene enrichment, it is identified both COVID-19 and lung cancer have a common pathway in EGFR tyrosine kinase inhibitor resistance, IL-17 signalling pathway, AGE-RAGE signalling pathway in diabetic complications, HIF-1 signalling pathway and pathways in cancer.

## Introduction

COVID-19 is a viral-based transmissible infection where the source of this disease outbreak is Severe Acute Respiratory Syndrome Coronavirus 2, also known as SARS-CoV-2 [1]. The outbreak of this virulent disease firstly was spotted in December 2019 originated from a small distinctive area of Wuhan, Hubei, China. Due to this issue, it has caused a continuous worldwide pandemic. The first confirmed case has been traced back to 17 November 2019 in Hubei [2]. As of 29^th^ January 2021, ~100 million confirmed cases globally ~2.1 million deaths for COVID-19 were reported. Human to a human transmission rate of SARS-CoV-2 infection has been seen at higher than that SARS epidemic observed in 2003. This rapid transmission of SARS-CoV-2 in human cases makes it extremely complex for prevention. The functional receptor which is necessary for the cellular entry of SARS-CoV-2 has been identified and discovered that angiotensin-converting enzyme 2 (ACE2) is playing a crucial role in cellular entry [3]. Since COVID-19 declared as a pandemic disease in early 2020, patients with a pre-existing health condition like cardiovascular disease, diabetes and cancer were detected as a potentially vulnerable population and having a high risk of death [4].

Lung cancer is the top cause of fatality among all cancer worldwide. Lung cancer can affect any people, especially people who smoke have the highest tendency of suffering from lung cancer, while lung cancer in people who have never smoked can also occur [5]. Recent research concerning potential risks of serious conditions related to each type of cancer-related to SARS-CoV-2 was conducted and discovered that COVID-19 and lung cancer both mutually related in terms of symptoms, diagnosis, and treatment [6]. For COVID-19, senior citizens and health-compromised individuals have a greater percentage of being sick. Specifically, individuals with lung cancer, individuals with lung disease, such as COPD (chronic obstructive pulmonary disease), and individuals in active cancer treatments may be susceptible to a more serious form of the infection [7]. These lung cancer patients or patients that undergo cancer therapies like chemotherapy are at high risks to suffer COVID-19 due to their weakened immune system. Reduced lung function and extreme infection in lung cancer patients may lead to the worse results in this subpopulation [8].

Coronavirus causes irritation and inflammation in the respiratory tracts lining as they travel themselves down our airway. In severe COVID-19 cases, the infection is carried on to both the lungs which cause serious inflammation and causes swelling which will lead our lungs to fill up with fluid and debris if it is worsening. In longer timeframe without quick treatment, the patients may have more serious pneumonia too. The lungs’ air sacs will be filled up with mucus, blood, and causes other immune cells to attempt to fend off the infection. This immunity mechanism requires a large amount of energy to execute and cause the body hard to intake oxygen. Due to these occurrences, many patients experienced breathing difficulties or shortness of breath as well as increase breathing rate. The higher mortality rate in COVID-19 patients has been correlated to the occurrence of the cytokine storm induced by SARS-CoV-2 virus. An extreme level of proinflammatory cytokines production leads to Acute respiratory distress syndrome (ARDS) [9].

Viral infections are the leading causes of acute and chronic infectious diseases worldwide. Identifying the mechanism of infection is the key challenge for drug discovery. Several studies have revealed how each virus has evolved to hijack the primary cellular pathways in humans and evade the innate immune response by altering important host cell factors. However, emerging, and re-emerging viruses like Dengue, Zika, Ebola and SARS-CoV-2 are shown to be the critical challenge for drug discovery in recent days [10]. These due to complex host-cell interaction mechanisms. Computational study of virus-host cell protein-protein interaction data is an efficient way to understand the molecular level viral infection mechanism [11, 12]. We hypothesized that linking a virus-host cell protein-protein interaction by gene network analysis would offer new insights for cancer patient treatments. Therefore, the main objective of this study is to implement protein to protein interaction in obtaining hub genes that contribute to both COVID-19 and lung cancer.

## Materials and methods

### COVID-19 data retrieval

COVID-19 curated gene sets downloaded from CTD database, DisGeNET and the study from Gordon et al [13]. All the unique genes were combined to construct the COVID-19 dataset of 5463 genes.

### Lung cancer data retrieval

Lung cancer genes were retrieved from CIViCmine database (http://bionlp.bcgsc.ca/civicmine/). The common overlapping genes of COVID-19 and lung cancer genes were obtained from venny tools (https://bioinfogp.cnb.csic.es/tools/venny/). Venny tool is a web-based method for creating Venn diagrams. It can be beneficial in genetic fields, where the specific set of genes expressed in one or even more data sources can be established. This web tools have been used to construct the Venn diagram showing overlapping data of COVID-19 and lung cancer.

### PPI interaction analysis

In this research, Cytoscape (https://cytoscape.org) is being used to construct and display the network of the protein-to-protein interaction of the intermediate genes between COVID-19 and lung cancer. The network was constructed by all the UniProt ID into the search query while using the STRING protein query. STRING (Search Tool for the Retrieval of Interacting Genes/Proteins) (https://string-db.org/) in molecular biology is a biological database as well as the web platform of documented and anticipated protein-protein interactions. Database STRING includes data from multiple sources from experimental evidence, strategies of numerical analysis, and selections of the published text. In this study, the ambiguous terms are resolved by setting the confidence score cutoff to 0.4 and their respective maximum additional interactors to zero to import the network.

### Determination of Hub Genes

All potential hub genes of a particular disease are discovered through the usage of cytoHubba, an application that is installed and executed in Cytoscape. CytoHubba was used to calculate the node’s scores. CytoHubba acts as an interface in a network to evaluate 11 scoring methods using the Maximal Clique Centrality (MCC) algorithm to show the top 10 ranked nodes, which may be the potential hub genes that cause COVID-19 infection in lung cancer patients. For gene enrichment purposes, the top 10 genes have been further analysed.

### Functional Enrichment of Hub Genes

For the hub genes enrichment further studies purposes, we have been using WebGestalt (WEB-based GEne SeT Analysis Toolkit) to do so. This software applications to combine functional enrichment analysis and knowledge visualization to handle, retrieve information, arrange, visualize and interpret large gene sets statistically [14]. In WebGestalt, all the top 10 genes are used in the analysis where two main analysis are studied which are their gene ontology and biological pathway which are displayed in charts.

## Result and discussion

The COVID-19 related curated genes are combined and retrieved from CTD, DisGeNET and Gordon study with a total of 5463 genes involved. The lung cancer genes were obtained from CIViCmine database with the total genes of 1750 genes while the 785-common overlapping COVID-19 and COPD genes were obtained from Venny tools which are presented in Venn Diagram as shown in Figure 1, respectively. The protein-to-protein interaction network of these common overlapping genes was created by using STRING in Cytoscape with the confidence score cutoff is set as 0.4 (Figure 3).

**FIGURE 1:**
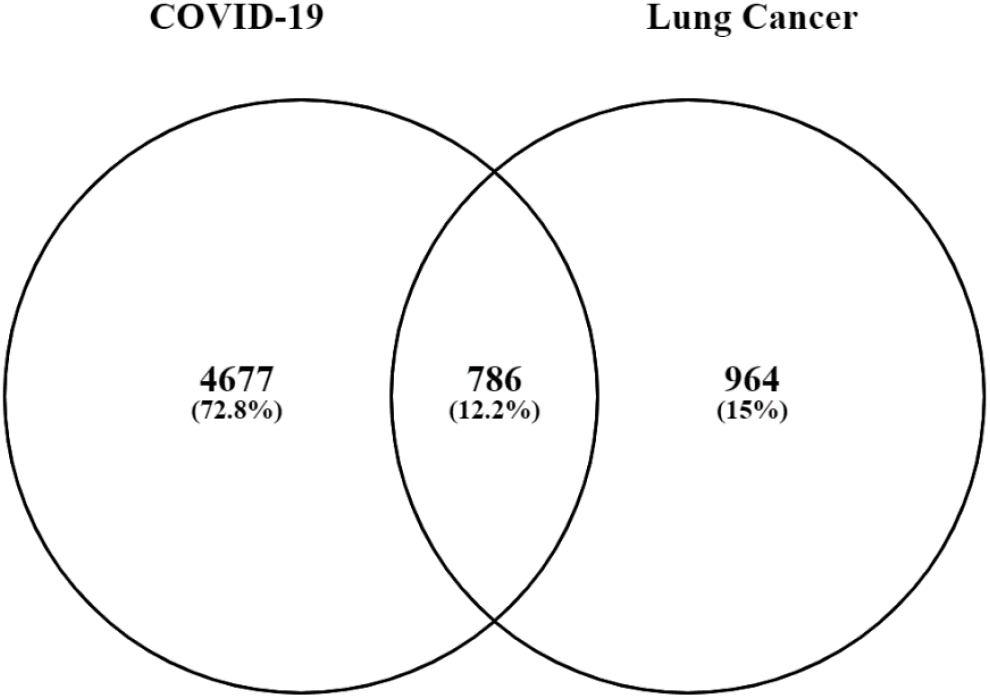
The intersection of Lung Cancer and COVID-19 genes.

CytoHubba was used to analyze the top 10 essential genes which were obtained using eleven different network analysis methods (Table 1). The rank of these nodes was obtained and differentiated according to a colour range which ranges from red to yellow. These hub genes, known by their gene symbol, are ALB, CXCL8, FGF2, IL6, INS, MMP2, MMP9, PTGS2, STAT3 and VEGFA shown in Figure 2 and their related genes which are SIRT2, FLT1, MYC, NRP1, KDR, CD63, PIK3CA, MET, ROS1, CEACAM1, CCND1, CD163 and ESR1. The gene enrichment analysis of these genes was made and displayed in bar graphs according to their biological processes, cellular component, a molecular function which is shown in Figure 4 as well as KEGG pathway shown in Figure 5.

**Figure 2:**
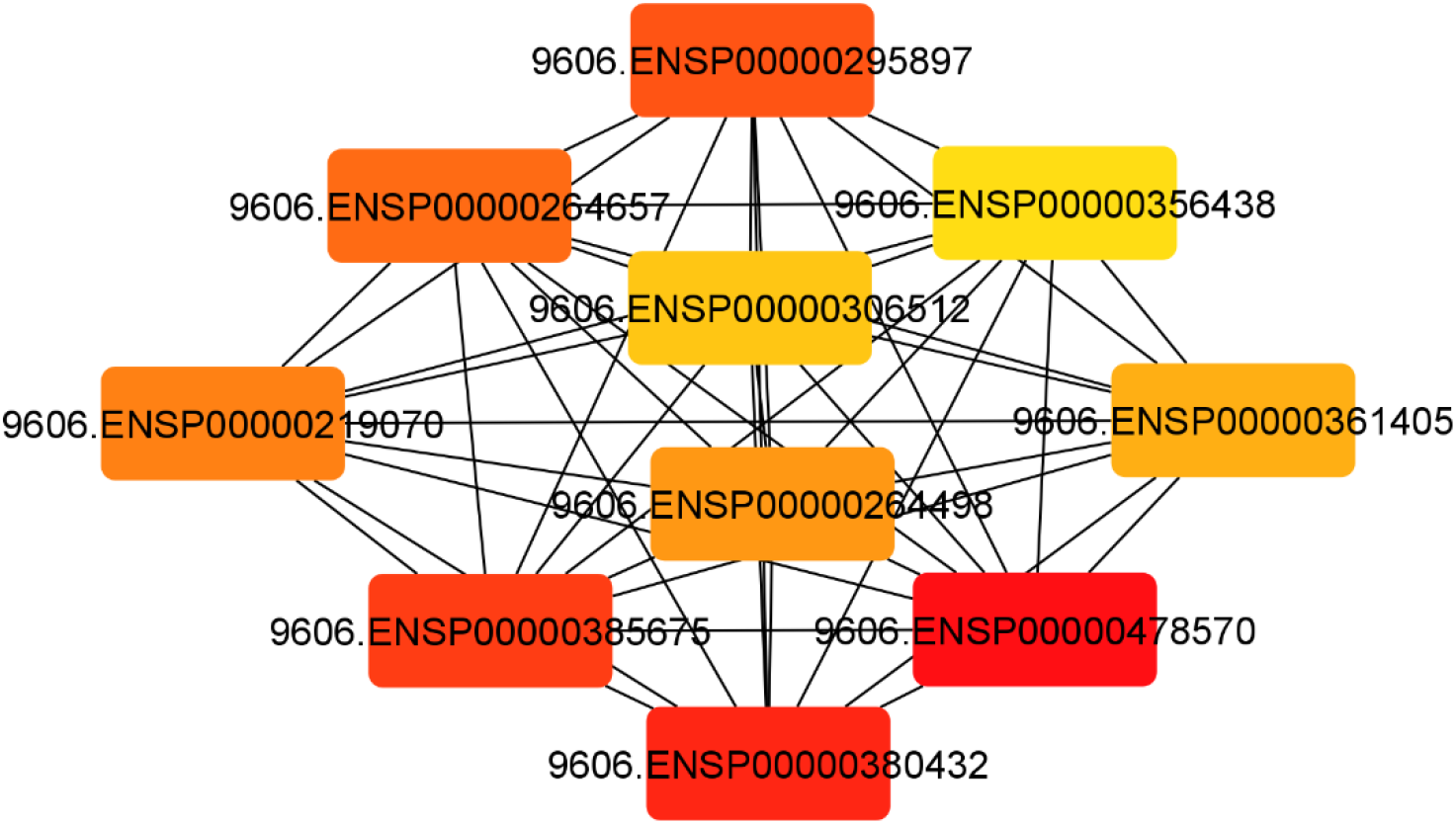
Protein to Protein Interaction Of Top 10 Genes.

**Figure 3:**
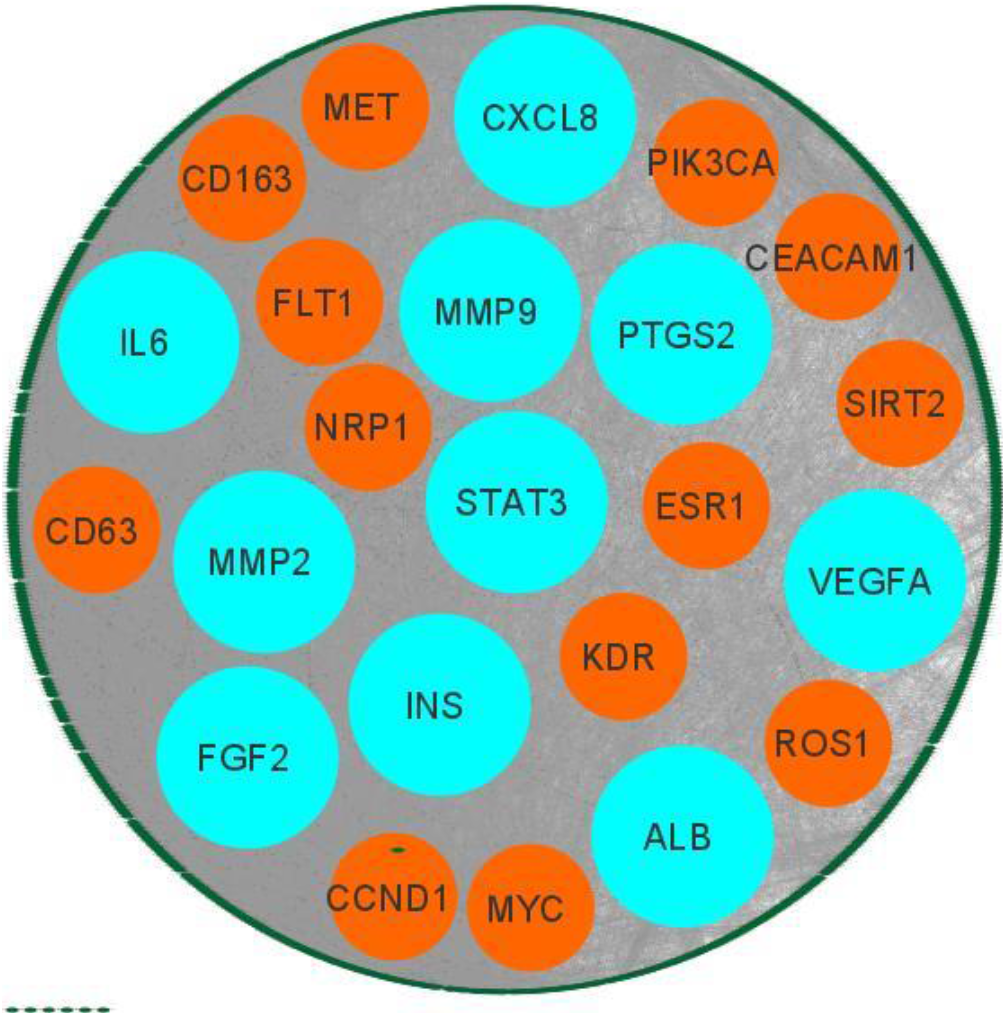
Shows the PPI Network.

**Figure 4:**
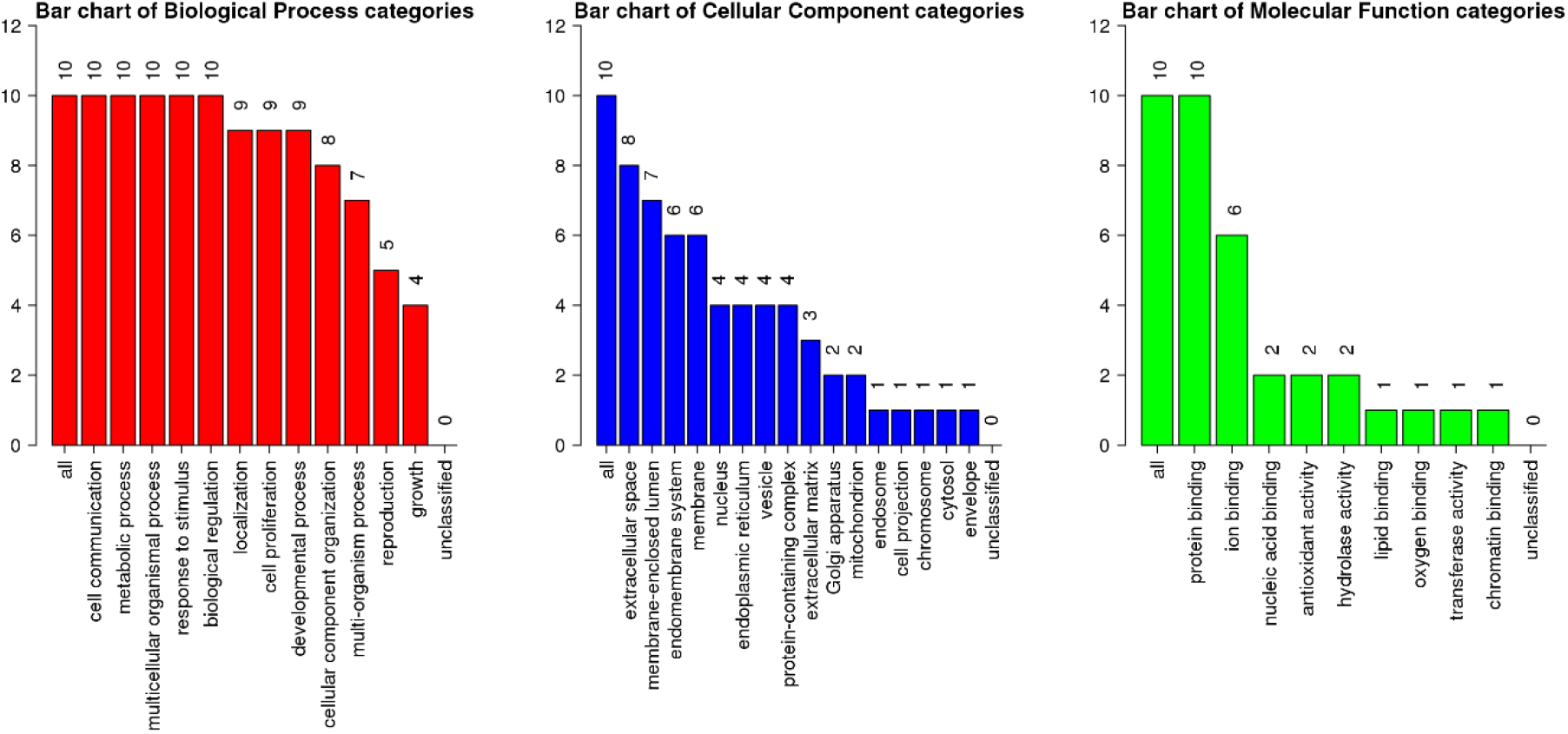
The Gene Enrichment of Top 10 Genes.

**Table 1:**
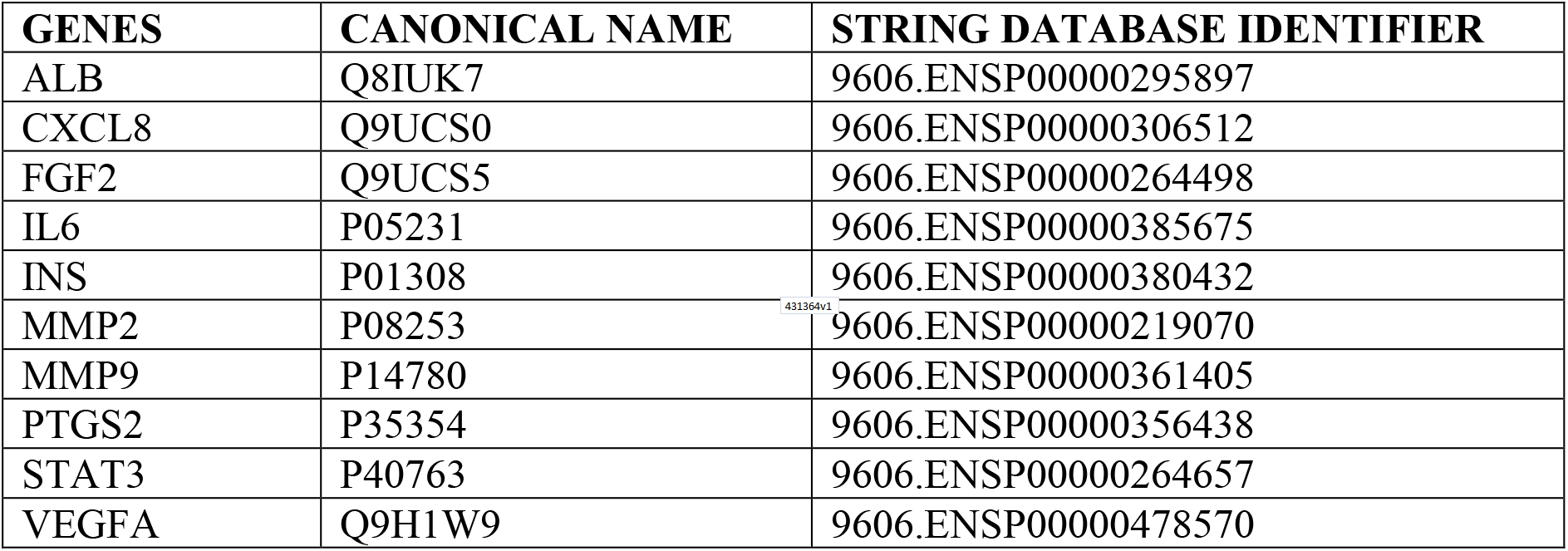
The List Of The Top 10 Genes.

According to 10 hub gene enrichment analysis (Figure 4), the first gene ontology analysis bar chart of biological process categories shows that all 10 hub genes are involved in cellular communication, metabolic process, multicellular organismal process, cellular response to a stimulus, and biological regulation. 9 of them are involved in localization, cell proliferation and developmental process, 8 hub genes involved in the cellular component organization, 7 genes involved in the multi-organism process, 5 genes involved in reproduction and lastly 4 genes involved in growth. Next hub gene enrichment analysis is on cellular components where 8 of the genes are found in extracellular space, 7 genes are found in the membrane-enclosed lumen, 6 genes are found in endomembrane system and membrane, 4 genes are found in the nucleus, endoplasmic reticulum, vesicle, and protein-containing complex, 3 genes are found in the extracellular matrix, 2 genes are found in Golgi apparatus and mitochondrion, and lastly 1 gene is found in the endosome, cell projection, chromosome, cytosol and envelope respectively. Last part of gene ontology analysis is on molecular function. All the hub genes are involved in protein binding, 6 genes in ion binding, 2 genes in nucleic acid binding, antioxidant activity and hydrolase activity. Lastly, 1 gene is involved in lipid, oxygen, chromatin binding and transferase activity respectively.

The five overlapping genes involved in positive regulation of smooth muscle cell proliferation (GO:0048661) are FGF2, IL6, MMP2, MMP9 and PTGS2. Next, six overlapping genes are involved in muscle cell proliferation (GO: 0033002) which are FGF2, IL6, MMP2, MMP9, PTGS2 and STAT3. CXCL8, FGF2, IL6, INS, MMP9, PTGS2, STAT3 and VEGFA are 8 overlapping genes that involved in positive regulation of cell migration (GO: 0030335), cell motility (GO: 2000147). Next gene set is positive regulation of cellular component movement (GO:0051272) consisting of 8 overlapping genes which are CXCL8, FGF2, IL6, INS, MMP9, PTGS2, STAT3 and VEGFA.

There are 8 overlapping genes involve in positive regulation of locomotion (GO:0040017) which are CXCL8, FGF2, IL6, INS, MMP9, PTGS2, STAT3 and VEGFA. Next, CXCL8, FGF2, IL6, MMP9, PTGS2, STAT3 and VEGFA are 7 overlapping genes that involve in angiogenesis (GO: 0001525. For cytokine-mediated signalling pathway (GO:0019221), there is 8 overlapping gene are CXCL8, FGF2, IL6, MMP2, MMP9, PTGS2, STAT3 and VEGFA. Meanwhile, under the regulation of cell migration gene set (GO: 0030334 the overlapping genes involved are CXCL8, FGF2, IL6, INS, MMP9, PTGS2, STAT3 and VEGFA. Lastly, there are 8 overlapping genes which are CXCL8, FGF2, IL6, INS, MMP9, PTGS2, STAT3 and VEGFA involved in the regulation of cell motility (GO:2000145).

KEGG pathway of all the top 10 genes (Figure 5) shows that they are greatly involved in bladder cancer (hsa05219), EGFR tyrosine kinase inhibitor resistance (hsa0152), AGE-RAGE signalling pathway in diabetic complications (hsa04933), IL-17 signalling pathway (hsa04657), HIF-1 signalling pathway (hsa04066), Kaposi sarcoma-associated herpesvirus infection (hsa05167), Hepatitis B (hsa05161), Proteoglycans in cancer (hsa05205), Human cytomegalovirus infection (hsa05163) as well as pathways in cancer (hsa05200).

**Figure 5:**
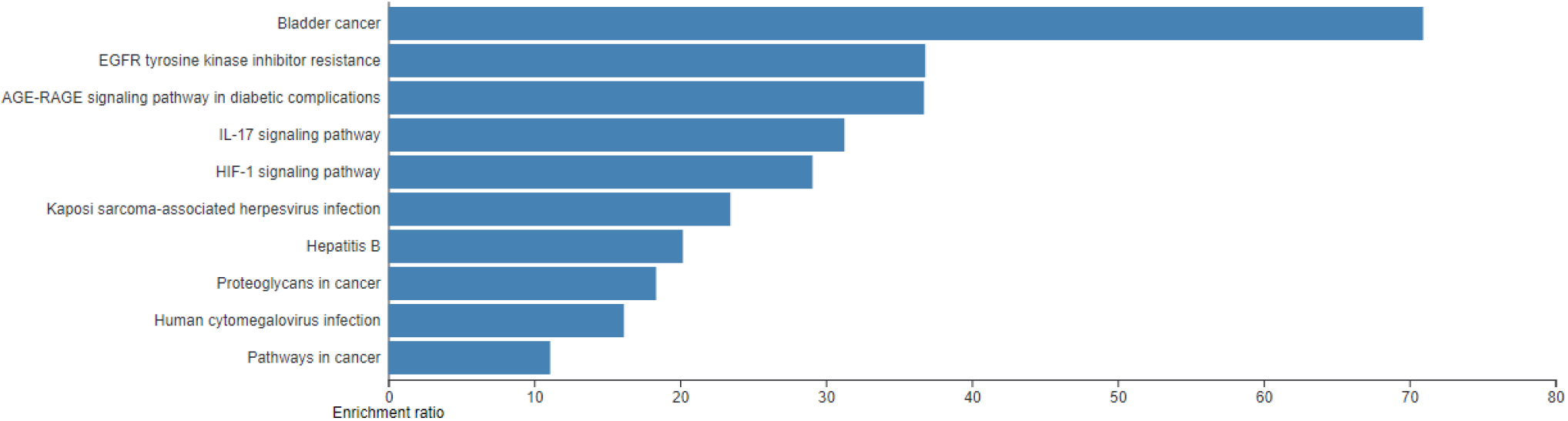
The pathway analysis of the top 10 genes.

For bladder cancer, there are 4 overlapping genes involved where are CXCL8, MMP2, MMP9 and VEGFA. EGFR tyrosine kinase inhibitor resistance consists of 4 overlapping genes involved which are FGF2, IL6, STAT3 and VEGTA. AGE-RAGE signalling pathway in diabetic complications contains 5 overlapping genes which are CXCL8, IL6, MMP2, STAT3 and VEGFA. An IL-17 signalling pathway consists of 4 overlapping genes which are CXCL8, IL6, MMP9 and PTGS2. A HIF-1 signalling pathway consists of 4 overlapping genes which are INS, IL6, STAT3 and VEGFA. Kaposi sarcoma-associated herpesvirus infection consists of 6 overlapping genes which are CXCL8, FGF2, IL6, PTGS2, STAT3 and VEGFA. Hepatitis B consists of 4 overlapping genes which are CXCL8, IL6, MMP9 and STAT3. Proteoglycans in cancer consist of 5 overlapping genes which are FGF2, MMP2, MMP9, STAT3, and VEGFA. Human cytomegalovirus infection consists of 5 overlapping genes CXCL8, IL6, PTGS2, STAT3 AND VEGFA. Lastly, pathway in cancer consists of 8 overlapping genes which are CXCL8, FGF2, IL6, MMP2, MMP9, PTGS2, STAT3 and VEGFA.

C-X-C motif chemokine ligand 8 (CXCL8) is the family of proteins is a subset of the CXC chemokine family and is a major inflammatory response mediator [15]. Mononuclear macrophages, neutrophils, eosinophils, T lymphocytes, epithelial cells and fibroblasts are secreted by IL-8. It acts as a chemotactic factor by directing the neutrophils to the infection site. This gene is responsible for creating inflammatory reactions in the lungs that lead to the excess fluid formation that cause breathing difficulties. Next, fibroblast growth factor 2 (FGF2) is a gene that encoded proteins which belongs to the family Fibroblast Growth Factor (FGF). The representatives of the FGF family attach heparin and have diverse mitogenic but also angiogenic activity. This protein has indeed been involved in various biochemical functions, like the development of the limb and nervous system, cell proliferation, and growth of tumours [16]. Interleukin 6 (IL6) is a gene that codes a cytokine that works in inflammation reaction and B cell growth and development [17]. In relation, it has been shown that the protein complex is an intracellular pyrogen able to induce fever in individuals with immune-mediated illnesses and diseases. It is understood that immune effector cells such as IL-6 and chemokines such as CXCL8 release during SARS-CoV-2 infection-causing ARDS by a cytokine storm. Therefore, patients with lung cancer are vulnerable to SARS-CoV-2 infection

Matrix Metallopeptidase 2 (MMP2) is the representative gene of the gene family matrix metalloproteinase (MMP), which are zinc-dependent enzymes incapable of slicing the extracellular matrix components and the proteins engaged in signal transduction [18]. Unlike other representatives of the MMP family, it can be activated upon on cell membrane. It is believed to be associated with several pathways involving functions in the nervous system, menstrual endometrial dissolution, vascularization control and cell proliferation. Matrix metallopeptidase 9 (MMP9) [19] protein plays an important role in the degradation of extracellular matrix in normal physiological processes, including embryonic development, reproduction and tissue remodelling, and illness mechanisms. MMP-9 is part of the protease family that degrades ECM proteins and is commonly studied in acute lung injury and chronic lung disease. MMP-9 is abundant in lung diseases characterised by tissue remodellings such as asthma, pulmonary fibrosis, and COPD, but poor in healthy lungs. MMP-9 released from neutrophils in acute lung injury causes swelling, alveolar-capillary barrier destruction, and further induces inflammatory cell migration and lung tissue degradation. The previous study suggests that MMP-9 could be an early predictor of respiratory failure in COVID-19 patients, emphasising the role of ECM remodelling and fibrosis in this condition. Treatment modalities for MMP-9 activity or neutrophil activation may be essential for COVID-19 affected lung disease patients. Prostaglandin-endoperoxide synthase 2 (PTGS2) is an important enzyme in prostaglandin biosynthesis and behaves either as dioxygenase but also peroxidase. There have been two PTGS isozymes which are constitutive PTGS1 and inducible PTGS2 which vary in their expression and tissue distribution regulation [20]. Specific stimulatory events regulate it, suggesting that it is responsible for the prostanoid biosynthesis involved in inflammation and mitogenesis. PTGS2 plays a pivotal role in inflammation, tissue damage, and tumorigenesis. Previous studies have suggested that FURIN and PTGS2 must play mechanistic roles in the formation of numerous signaling factors that promote a malignant phenotype in non-small cell lung cancer (NSCLC) [21]. Dysregulation of FURIN leads to enhanced bioavailability of hormones and growth factors that promote tumour progression. Furin is enriched in the Golgi apparatus, where it functions to cleave other proteins into their mature/active forms the spike protein of SARS-CoV-2 to become fully functional.

Signal transducer and activator of transcription 3 (STAT3) is an encoded protein part of the STAT family of proteins. STAT family members are phosphorylated by the receptor-associated kinases in response to cytokines and growth factors and then form homo- or heterodimers that translocate to the nucleus of the cell which they serve like transcription activators [22]. In the regulation of host immune and inflammatory responses as well as in the pathogenesis of many cancers STAT3 plays a major role. In a variety of viral infections, several studies have documented differential regulation of STAT3. Transcription transducer and activator 1 (STAT1) dysfunction and compensatory hyperactivation of STAT3 are caused by viral SARS-CoV-2 components. The initial and mild phases of the SARS-CoV-2 infected cells and alveolar macrophages are limited to type 2. When infected, the SARS-CoV-2 infestations triggered by STAT3, include IP-10 (CXCL10), facilitate the secretion of MMP-9 by alveolar cell types 2, including SARS-CoV-2. PAI-1 was found to be overexpressed in human NSCLCs. In severe cases of COVID-19, there is a common escalating cycle of STAT3 and PAI-1 activation that is shared among diverse disease manifestations and leads to catastrophic consequences [23].

Vascular endothelial growth factor A (VEGFA) belongs to both the PDGF and VEGF family of growth factors. It produces a protein that binds heparin, which occurs as a homodimer connected to the disulphide [24]. This growth factor stimulates the differentiation and proliferation of endothelial vascular cells and is important in both physiological angiogenesis as well as pathology. Clinical results showing increased VEGF-A level in COVID-19 patients with the bronchial alveolar lavage fluid. Interestingly, an analysis of gene expression in patients with COVID-19 showed upregulation of NRP1 and NRP2 in lung tissue. In the previous study, it is identified that SARS-CoV-2 Spike protein hijacks NRP-1 signalling to enhance VEGF-A mediated pain. This increases the potential for the pain to be diffused directly as an early symptom of COVID-19 by the SARS-CoV-2 spike protein.

Albumin (ALB) is a gene that code a most plentiful protein within the bloodstream of humans. This protein works in regulating osmotic potential in the blood plasma as well as behaves as a carrier protein for a wide range of endogenous molecules including hormones, fatty acids and metabolites, and exogenous drugs [25]. Hypoalbuminemia has now been reported in patients with severe disease seeking help in the emergency room because of COVID-19 infection. albumin synthesis in the hepatocyte is downregulated at a pre translational level by the direct interaction of the major acute-phase cytokines which are released into the circulation during the cytokine "storm" induced by the viral effects on the lungs [26].

Low serum albumin levels have now been found to be an important predictor of progression to severe disease and increased mortality in hospitalised SARS-CoV-2 positive patients of older age [26]. Leaking of albumin into the interstitium increases the colloid oncotic pressure in this space, and may worsen conditions, such as acute respiratory distress syndrome. Trauma, disease, growth, etc. are inflammatory conditions that are associated with hypoalbuminemia. More severe inflammation is associated with progressively lower serum albumin levels, although the strength of the correlation has only sparsely been investigated. There is a highly significant correlation between serum albumin level and mortality risk when stratified for age and gender [27].

Insulin (INS) is the gene that provides instructions for making insulin, a peptide hormone which plays a significant role in carbohydrate regulation or even lipid metabolism [28]. Insulin was also shown to increase ACE2 expression by attenuating disintegrin and metalloprotease (ADAM-17) influence. In recent years, it has become apparent that ADAM-17 protease can alter several non-matrix substrates such as cytokines (e.g., TNF-α), cytokine receptors (e.g., IL-6R and TNF-R), ErbB ligands (e.g., TGF-α and amphiregulin) and adhesion proteins (e.g. L-selectin and ICAM-1) [29]. ADAM17 participates in ACE2 ectodomain shedding, TMPRSS2 induces ACE2 intracellular cleavage, and there was competition for ACE2 processing between ADAM17 and TMPRSS2. Recently, it has been stated that SARS-CoV-2 uses TMPRSS2 protease for S-protein processing and that TMPRSS2 inhibitors have blocked SARS-S cell entry [30].

Many studies have shown that IL-17 promotes tumour angiogenesis and cell proliferation directly or indirectly and inhibits apoptosis by triggering inflammatory signalling pathways. IL-17 thus leads to lung cancer metastasis and development [31]. Patients with exuberant T-cell activation, and Th17 cellular infiltration, resulting in a rise in inflammatory cytokines such as interleukin-17A (IL-17A) (also referred to as IL-17) after SARS-CoV-2 infection. IL-17 is an important and predominant mediator of pulmonary inflammation among many cytokines involved in the storm. IL-17’s pro-inflammatory properties also make it essential for inflammatory and immunopathological mediators.

## Conclusion

Protein-protein interactions and network analysis are important in understanding protein functions and behaviour as well as analysis studies. Throughout this study, we have identified the 10 hub genes of COVID-19 and Lung cancer. The results indicate that most of these genes encode for binding proteins that are involved in biological pathways. The identified hub genes through network analysis can be used to further study the relation of COVID-19 and lung cancer and treatment discoveries.

